# Concurrent Genome and Epigenome Editing by CRISPR-Mediated Sequence Replacement

**DOI:** 10.1101/675447

**Authors:** Jes Alexander, Gregory M. Findlay, Martin Kircher, Jay Shendure

## Abstract

Recent advances in genome editing have facilitated the direct manipulation of not only the genome, but also the epigenome. Genome editing is typically performed by introducing a single CRISPR/Cas9-mediated double stranded break (DSB), followed by NHEJ or HDR mediated repair. Epigenome editing, and in particular methylation of CpG dinucleotides, can be performed using catalytically inactive Cas9 (dCas) fused to a methyltransferase domain. However, for investigations of the role of methylation in gene silencing, studies based on dCas9-methyltransferase have limited resolution and are potentially confounded by the effects of binding of the fusion protein. As an alternative strategy for epigenome editing, we tested CRISPR/Cas9 dual cutting of the genome in the presence of *in vitro* methylated exogenous DNA, *i.e.* to drive replacement of the DNA sequence intervening the dual cuts via NHEJ. In a proof-of-concept at the *HPRT1* promoter, successful replacement events with heavily methylated alleles of a CpG island resulted in functional silencing of the *HPRT1* gene. Although still limited in efficiency, our study demonstrates concurrent epigenome and genome editing in a single event, and opens the door to investigations of the functional consequences of methylation patterns at single CpG dinucleotide resolution. Our results furthermore support the conclusion that promoter methylation is sufficient to functionally silence gene expression.

## Introduction

Mammalian genome editing has become much more straightforward with the discovery of CRISPR. Conventional genome editing with CRISPR uses the endonuclease Cas9 to cut the genome at a programmed location which is followed by endogenous DNA repair (Sander and Keith Joung 2014). The targeting of the Cas9 cut is programmed by a guide RNA which has homology to the sequence that will be cut by Cas9. DNA repair occurs through two main pathways: homology directed repair (HDR) and non-homologous end joining (NHEJ). HDR-mediated genome editing requires an exogenous DNA repair template bearing homology arms that are used in homologous recombination of the template with the genome, resulting in a precise change at the position of the programmed cut. In contrast, NHEJ-mediated genome editing simply involves religating the broken ends, but this occasionally results in small insertions or deletions, *i.e.* an imprecise change at the position of the programmed cut. However, if an exogenous DNA template is provided, it can be inserted at the location of the programmed cut by NHEJ-mediated ligation (Tsai et al. 2015). If dual cuts are programmed nearby to one another, NHEJ-mediated ligation at both double-stranded breaks can result in the replacement of the intervening sequence with an exogenous DNA template (Geisinger et al. 2016).

Although the ability to edit the base sequence of the genome is very useful, much of the information that carries cell type-specific properties, such as gene expression, is encoded at an epigenetic level. CpG island methylation is one such layer of epigenetic regulation (Dirk Schübeler 2015; Jones 2012). CpG dinucleotide methylation is important in both normal development as well as disease, but the mechanisms by which it contributes to the regulation or dysregulation of gene expression remains poorly understood (Bergman and Cedar 2013; Jin and Liu 2018).

Editing of DNA methylation has previously been demonstrated by two approaches. In a first approach based on site-specific recombinases such as Cre-*loxP, loxP* sites are integrated into the genome at a locus of interest; an *in vitro* methylated plasmid with *loxP* sites is then transfected and Cre recombinase expressed; this drives recombination of the *in vitro* methylated DNA into the genome at the locus of interest (M. C. Lorincz, Schübeler, and Groudine 2001; Matthew C. Lorincz et al. 2002; D. Schübeler et al. 2000). This approach is highly efficient, but major drawbacks include that the *loxP* sites must be engineered into the genome first, and these sites remain in the genome even after recombination.

A second, recently demonstrated approach uses a catalytically inactive Cas9 as a targeting domain fused to a DNA methyltransferase domain for methylation of CpG dinucleotides (Liu et al. 2016; Amabile et al. 2016; Vojta et al. 2016; McDonald et al. 2016; Stepper et al. 2017; Lei et al. 2017; Xiong et al. 2017; Adli 2018). This approach has lower efficiency and results in methylation of multiple CpGs surrounding the target site, requiring multiple guides if the goal is to methylate a region. In the case of a CpG island, guide design can be complicated by low sequence complexity and targeting ambiguities. For investigations of the functional consequences of methylation, a limitation of this approach is that fails to discriminate between the consequences of the binding of the fusion protein vs. methylation itself.

We wondered whether it would instead be possible to achieve genome editing with respect to CpG methylation by using CRISPR/Cas9 to introduce DSBs at two nearby locations, followed by replacement of the intervening segment with a transfected, *in vitro* methylated version of the same sequence via NHEJ-mediated ligation (**Fig. 1a**). This strategy has the potential to enable methylation of an entire CpG island (hundreds to thousands of bases) with only two guides. It would also facilitate the introduction of precise, complex patterns of methylation or even of other DNA modifications. Finally, it represents concurrent genome and epigenome editing (*i.e.* if the exogenous, methylated segment differed in its base sequence from the endogenous segment). To test this approach, we targeted methylation to the CpG island of *HPRT1* in human Hap1 cells (Gasperini et al. 2017). *HPRT1* is a housekeeping gene with the special property that loss of its expression, whether by silencing or mutation, results in resistance to 6-thioguanine (6-TG), a chemotherapeutic purine analog. The Hap1 cell line is haploid which means that modification of only a single copy of the *HPRT1* locus is required to observe this phenotype.

**Figure 1.**
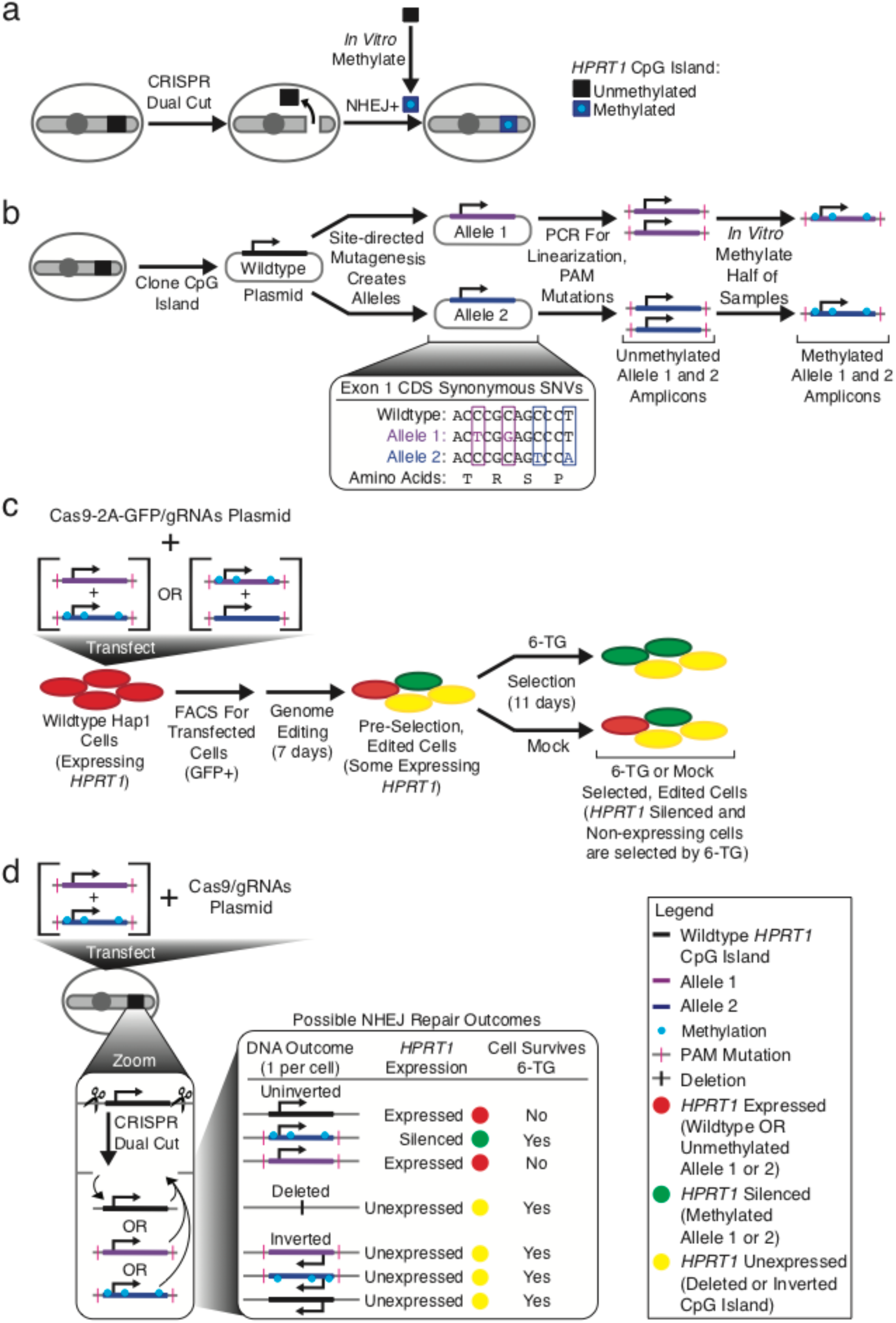
Experimental Design. **(a)** Overview of the experimental approach showing CRISPR dual cuts for removing and replacing the *HPRT1* CpG island with an *in vitro* methylated DNA sequence through NHEJ-mediated repair. **(b)** The *HPRT1* CpG island was cloned and synonymous coding SNVs were introduced to create two distinguishable alleles (blue and purple). Cloned CpG island alleles were PCR amplified for linearization and to incorporate PAM mutations. Portions of the resulting amplicons were *in vitro* methylated (cyan) with M.SssI. **(c)** For each replicate, the methylated version of one allele amplicon and the unmethylated version of the other allele amplicon, together with plasmids bearing Cas9-2A-GFP and two gRNAs, were co-transfected into Hap1 cells. In one plate of Hap1 cells, allele 1 was methylated and allele 2 was not, and in a parallel experiment, allele 2 was methylated and allele 1 was not. Transfected cells were sorted by FACS and replated for genome editing. Edited cells were then either selected with 6-TG, which will select for cells that do not express *HPRT1*, or mock selected with DMSO. Cells were harvested before and after selection, DNA was extracted, and the relevant regions PCR amplified and sequenced. The alleles allow tracking of the inserted methylated vs. unmethylated CpG island amplicons without requiring bisulfite conversion. The relative frequencies of the methylated and unmethylated alleles were calculated and compared between the 6-TG selected, mock selected, and pre-selection cells. **(d)** Potential outcomes of genome editing are shown for a hypothetical single cell from a single replicate. After a CRISPR dual cut, the possible outcomes at the DNA level are a deletion of the CpG Island, re-insertion of the original wild-type CpG island that was cut out, or insertion of the methylated or unmethylated alleles that were transfected in. Inserted CpG islands can be inverted or uninverted. *HPRT1* will be expressed if either the original wild-type or the unmethylated allele is inserted, but will no longer be expressed if a deletion or inversion occurs. Insertion of the uninverted, methylated allele should result in methylation-induced silencing. Finally, cells are expected to survive 6-TG selection if they no longer express *HPRT1*, which can be a consequence of methylation-induced silencing, deletion of the CpG island, or inversion of the CpG island. Therefore, on sequencing after 6-TG selection, if the methylated allele is inserted, we predict that its relative frequency will be increased as compared to the unmethylated allele.

## Results

We attempted to replace the *HPRT1* CpG island with *in vitro* methylated DNA using CRISPR-mediated NHEJ (**Fig. 1a**). To this end, the *HPRT1* CpG island, which overlaps with the first exon of *HPRT1* including a portion of the ORF, was cloned from human genomic DNA (**Fig. 1b**). Two synonymous SNVs were introduced into the coding sequence of the first exon in the cloned plasmid construct to generate a first allele that was distinguishable from the wild-type CpG island sequence. From the starting construct, a second allele was also created by two synonymous SNVs at different positions than used for the first allele. Since the positions used for the synonymous SNVs in the two alleles were different, the alleles were distinguishable from one other as well as the wild-type sequence. The CpG island alleles were PCR amplified to linearize them and then *in vitro* methylated with the enzyme M.SssI. Through the primers used for this PCR, mutations were introduced to the locations corresponding to the PAM site of the intended guide RNA targets, in order to reduce the probability of re-cutting by Cas9 after any successful insertion events (**Fig. 1b**; **Supp. Fig. 1**).

Methylated allele 1 and unmethylated allele 2 amplicons, along with plasmids directing expression of Cas9-2A-GFP and guide RNAs targeting the ends of the *HPRT1* CpG island, were co-transfected into a single plate of Hap1 cells. The reciprocal experiment, *i.e.* using a methylated version of allele 2 and an unmethylated version of allele 1, was performed in parallel, as a form of replication as well as to control for any effects of the synonymous mutations (**Fig. 1c**). Both the primary and reciprocal experiment were performed in triplicate. A key point is that with this experimental design, the alleles allow one to infer whether the methylated or unmethylated amplicon was inserted, without requiring bisulfite conversion prior to sequencing.

At 48 hours after transfection, >100,000 GFP positive cells were sorted by FACS and put back into culture for 7 days. GFP positivity indicates that these cells were transfected successfully. At this point, half of the cells from each plate were harvested (“pre-selection” in **Fig. 1c**) and the remaining half of the cells were split to two dishes. To one dish, 6-TG was added as a selection agent (“6-TG selected” in **Fig. 1c**) and to the other dish, DMSO was added as a vehicle control (“mock selected” in **Fig. 1c**). After 11 days, cells were harvested, genomic DNA was extracted, and the *HPRT1* CpG island was PCR amplified and sequenced.

Based on sequencing, the relative frequencies of methylated and unmethylated alleles were calculated and compared between pre-selection, mock selected, and 6-TG selected samples. These frequencies are dependent on the outcomes of genome editing, as expected outcomes are expected to lead to survival or death under 6-TG selection (**Fig. 1d**). Possible editing outcomes include deletion of the intervening segment, re-insertion of the original wild-type CpG island, or insertion of the transfected methylated or unmethylated allele. Additionally, the wild-type CpG island or methylated or unmethylated alleles can potentially be inserted in the original or an inverted orientation. Since Hap1 cells are haploid, only one of these editing outcomes is expected per cell. Insertion of the methylated allele in the uninverted orientation might be expected to result in methylation-induced silencing of *HPRT1*, while a deletion or any inversion would result in loss of expression. Cells with silencing or loss of expression of *HPRT1* are expected to survive 6-TG selection, while those with expression are expected to be strongly selected against.

We first sequenced the allele-defining SNVs and the surrounding portion of exon 1 using short-read Illumina sequencing. For this, a nested PCR approach was employed with one outer nest PCR primer upstream of the 5’ cut site and one between the cut sites (**Supp. Fig. 2**). The inner nest amplified a 44-bp region including the allele-defining SNVs in the exon 1 CDS and a small portion of the promoter, the amplicons from which were sequenced. This nested approach prevented amplification and sequencing of any random integrations at other positions in the genome, on-target inserts that were inverted, and deletions.

**Figure 2.**
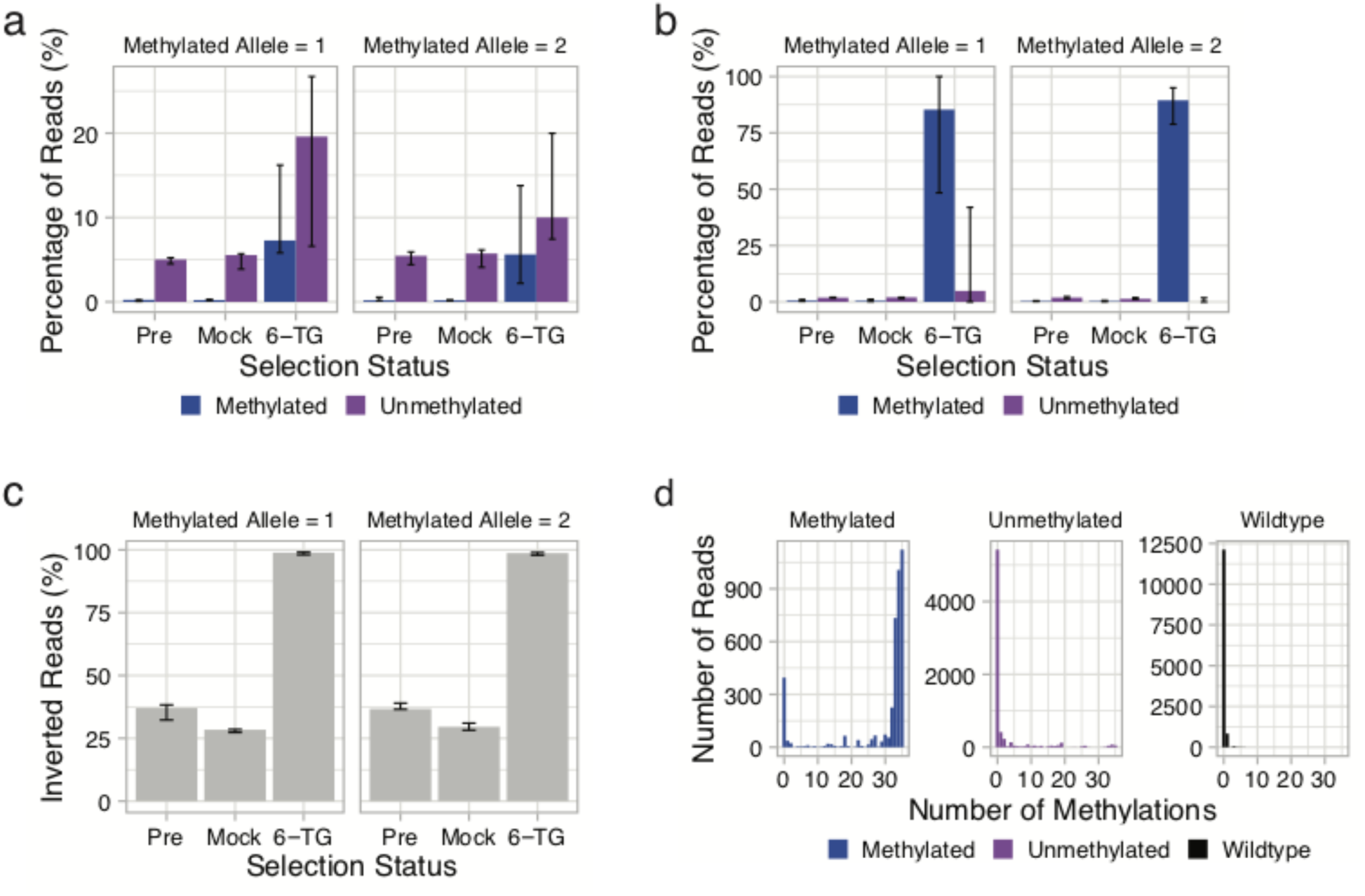
Methylation of the *HPRT1* CpG island by CRISPR-mediated sequence replacement results in *HPRT1* silencing. **(a)** Percentages of Illumina sequencing reads assigned to methylated and unmethylated inserted alleles by SNVs, grouped by selection status (Pre = pre-selection; Mock = mock selection; 6-TG = 6-TG selection). Although both are enriched, methylated inserted alleles are more enriched than unmethylated inserted alleles after 6-TG selection. Wild-type sequences are not shown, but are included in percentages. The first panel shows the experiment where allele 1 was methylated and allele 2 unmethylated; the second panel shows the reciprocal experiment. Error bars show the range. **(b)** Percentages of PacBio sequencing reads assigned to “exact matching” methylated and unmethylated inserted alleles by SNVs, grouped by selection status (Pre = pre-selection; Mock = mock selection; 6-TG = 6-TG selection). Methylated inserted alleles, but not unmethylated inserted alleles, are strongly enriched by selection. Sequences were only counted if they were in the forward orientation and exactly matched the promoter, exon 1, splice donor, PAM mutation, and one of three sets of allele-defining SNVs (wild-type, allele 1 or allele 2). Wild-type sequences are not shown, but are included in percentages. Error bars show the range. (**c**) Percentages of PacBio sequencing reads assigned to reverse/inverted orientation and grouped by selection status. Deletion events, as well as sequences not meeting the “exact matching” criteria defined above, were not counted. Forward oriented sequences are not shown, but are included in percentages. The clear pattern is that inverted sequences predominate after 6-TG selection. **(d)** Observed number of methylated sites upon bisulfite sequencing of methylated, unmethylated or wild-type allele of CpG island. Region contains 35 CpG dinucleotides. Reads are assigned to *in vitro* methylated or unmethylated alleles, or to unedited wild-type sequence based on synonymous SNVs. *In vitro* methylated alleles remain heavily methylated while unmethylated alleles and unedited sequences remain predominantly unmethylated.

Because these other outcomes are excluded by the nested PCR approach, our expectations were as follows: If the methylated allele is inserted, 6-TG selection should result in an increase in the frequency of the methylated allele compared to the unmethylated allele (quantifiable by sequencing of the allele-defining SNVs). In contrast, in the pre-selection and mock selection samples, no difference in the frequency of methylated and unmethylated alleles is predicted.

We quantified the frequencies of the inserted methylated and unmethylated alleles, both pre-selection as well as after 6-TG and mock selection (**Fig. 2a**). A first observation is that even pre-selection, the proportion of methylated alleles that are inserted is very low (mean 0.24%). In contrast, the proportion of unmethylated alleles inserted is modest but consistent (mean 5.1%). This suggests that NHEJ-mediated insertion of methylated alleles is markedly less efficient than that of unmethylated alleles. For both methylated and unmethylated alleles, the proportions after mock selection were largely unchanged. Surprisingly, the effect of 6-TG selection was to increase the percentage of *both* the inserted methylated and unmethylated alleles, relative to the wild-type allele. However, the fold-change for 6-TG selection over mock selection on the methylated allele was much greater than that of the unmethylated allele, suggesting an enrichment for the methylated allele, which is consistent with methylation-induced silencing of *HPRT1* (mean fold-change for methylated vs. unmethylated: 41.0 vs. 3.0; log-transformed, paired t-test: p ≈ 0.002).

We speculated that the unexpected increase in inserted, unmethylated alleles upon selection might result from loss of expression due to repair-induced indels at the ends of the of the CpG island insert (in the first intron or 5’ UTR), or alternatively from mutations in the promoter, exon 1 coding sequence, or splice-donor in the CpG island insert introduced by PCR. To test this, we amplified a ~2 kb region including the entire CpG island, with primers positioned ~700 base pairs (bp) upstream of one cut site and ~165 bp downstream of the other cut site (**Supp. Fig. 3**). We sequenced these amplicons using Pacific Biosciences (PacBio) instruments (**Methods**).

**Figure 3.**
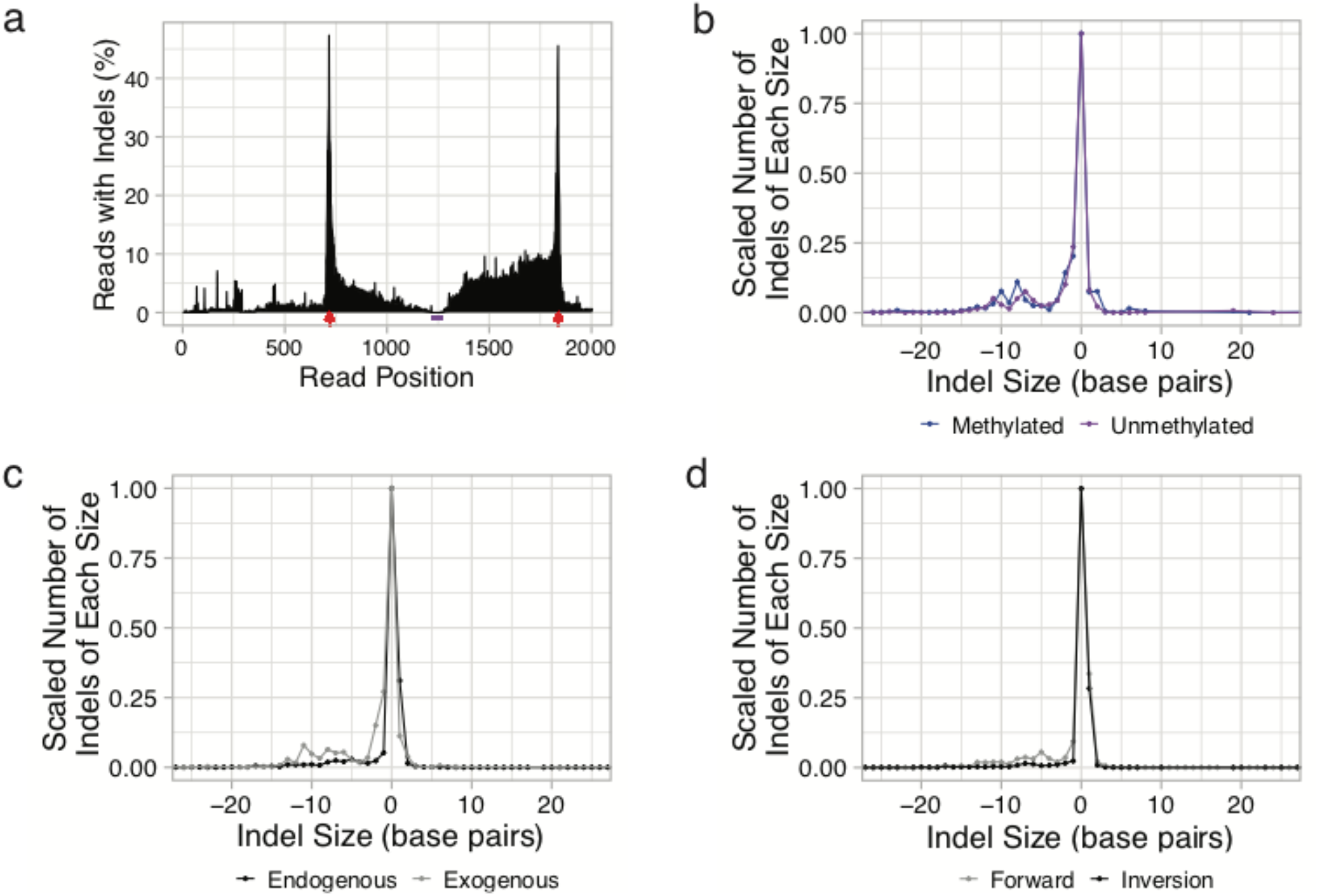
The positional and size distribution of indels, in relation to methylation status, insertion type and orientation. **(a)** Percentage of reads with an indel at positions along the PacBio sequenced region. The same subset of reads used in Fig. 2a are included here (6-TG selected, both methylated and unmethylated, perfect match on the allele-defining SNVs and the surrounding portion of exon 1). Red arrowheads indicate the CRISPR/Cas9 cut sites. The purple bar marks the region of exon 1 surrounding the allele-defining SNVs. The distribution of indels is highest at the CRISPR/Cas9 cut sites, but many reads have indels within the CpG island as well. **(b)** Indel distributions of methylated (blue) or unmethylated (purple) alleles. **(c)** Indel distributions from events involving exogenous inserts (gray) or endogenous inserts (black) **(d)** Indel distributions from forward-oriented wild-type sequences (gray) or inverted wild-type sequences (black). The numbers of indels (y-axis) were scaled so that the maximum number for any indel size (x-axis) for a given distribution was one to allow easier comparison between distributions. Negative numbers for indel size represent deletions, and positive numbers represent insertions.

Circular consensus sequence (CCS) calling was performed to a mean CCS accuracy of 99.4%. In contrast with our Illumina-based sequencing, this approach is expected to recover not only forward-oriented alleles, but also inversions and deletions. However, for this first analysis, sequences were only counted if they were in the forward orientation and moreover exactly matched the promoter, exon 1, splice donor, PAM mutation, and one of three sets of allele-defining SNVs (wild-type, allele 1 or allele 2), *i.e.* to exclude sequences containing PCR errors or repair-induced indels. In contrast with results presented in **Fig. 2a**, after 6-TG selection, we observed markedly stronger selection for methylated inserted alleles than for unmethylated inserted alleles (mean 82.8% vs. 8.1%; arcsine square root transformed, paired t-test: p ≈ 0.005) (**Fig. 2b**). However, as illustrated by the pre-selection and mock selection experiments, the proportion of inserted methylated and unmethylated alleles remained very low in the absence of 6-TG. As the main difference between the Illumina and PacBio-based analyses related to the latter requiring “exact matching” across the entire region, these results are consistent with the view that repair-induced indels and/or PCR errors result in a subset of forward-oriented, unmethylated inserts being positively selected. We return to this topic further below.

We also examined other sequences in the PacBio data, *i.e.* sequences other than those exactly aligning to forward-oriented wild-type or inserted alleles. For example, one prediction is that 6-TG should also select for alleles inserted in the inverted orientation, regardless if it is the wild-type sequence or one of the exogenous inserts. To investigate this, we tabulated sequences that exactly matched the promoter, exon 1, splice donor, PAM mutation, and any of three sets of allele-defining SNVs (wild-type, allele 1 or allele 2), in either orientation. Collapsing all alleles in each orientation, we observe that the proportion of forward-oriented alleles was modestly higher in both the pre-selection and mock selection samples (mean 63.4% and 71.1% forward oriented, respectively). However, after 6-TG selection, the overwhelming majority of sequences were in the reverse/inverted orientation (mean 98.6% reverse oriented) (**Fig. 2c**). This confirms that 6-TG selection was nearly complete, particularly as the forward-oriented sequences observed after 6-TG selection were dominated by the methylated, inserted alleles (**Fig. 2b**).

Although we observe that the forward oriented, methylated allele is strongly selected for by 6-TG, we sought to confirm that its *in vitro* methylation is maintained after transfection and insertion, and thus might plausibly cause silencing of *HPRT1* and the consequent selection. We therefore performed bisulfite sequencing on a region of the CpG island including the allele-defining SNVs and 35 surrounding CpGs (**Supp. Fig 4**). We observe that the *in vitro* methylated allele remained heavily methylated in the pre-selection, mock selection, and 6-TG selection samples, whereas the unmethylated allele and the wild-type sequence remained predominantly unmethylated in all samples (**Fig. 2d**).

Although our strategy remains challenged by the much higher rates of deletion or inversion over insertion of the methylated inserts, our observations nonetheless support the conclusions that: (a) we successfully used CRISPR/NHEJ to replace the *HPRT1* CpG island with an *in vitro* methylated allele; (b) this methylation was maintained after insertion to the genome, at least over the course of our 11 day experiment; and (c) this methylation was sufficient to functionally silence the *HPRT1* gene.

We further analyzed the PacBio sequencing data to explore the indel patterns associated with CRISPR/NHEJ-mediated sequence replacement. First, we revisited the question of why, in the Illumina short-read sequencing, 6-TG selection resulted in enrichment of both methylated and unmethylated alleles rather than just methylated alleles (**Fig. 2a,b**). As discussed above, comparison of the Illumina short-read sequencing and PacBio sequencing data suggested that indels in functional regions of the CpG island insert, *i.e.* the 5’ UTR, promoter, exon 1, or splice-donor sequences, might cause loss of expression of *HPRT1*, resulting in selection of these indel-bearing, unmethylated sequences by 6-TG. We formally addressed the question by analyzing the distribution of indels across the region subjected to PacBio sequencing (**Fig. 3a**). Both methylated and unmethylated allele sequences selected by 6-TG, with a perfect match of the allele-defining SNVs and the surrounding region of exon 1, were included. This was the same inclusion criteria used for the Illumina short-read sequencing to facilitate comparison. As expected, the indel distribution had peaks at both CRISPR/Cas9 cut sites (**Fig. 3a**). Notably, many indels extended from the flanking CRISPR/Cas9 cut sites into the interior of the CpG island encompassing functional regions involved in *HPRT1* expression. Such indels are predicted to result in loss of expression of *HPRT1*. Since these regions were not visible to Illumina short-read sequencing, indel-containing alleles were included in the results shown in **Fig. 2a**, but were excluded by our sequencing matching requirements with PacBio for the results shown in **Fig. 2b**. Overall, we conclude that any modest enrichment of unmethylated alleles after 6-TG selection was likely due to these alleles containing indels extending into functional regions of the CpG island.

We next examined the effects of methylation on indel patterns in CRISPR/NHEJ-mediated sequence replacement. We started by asking whether there are differences in the rates of insertion of methylated vs. unmethylated alleles. Combining alleles and observations in both orientations, we observed that the unmethylated allele was consistently incorporated more frequently than the methylated allele (0.40% methylated vs. 1.54% unmethylated in pre-selection; 0.35% methylated vs. 1.26% unmethylated in mock selection). These differences were consistent between forward and reverse oriented events. There have been reports that some double strand breaks in methylated DNA are repaired differently than those in unmethylated DNA (Thongsroy et al. 2013; Kongruttanachok et al. 2010). While we cannot rule out an artifact of our experimental setup, it is possible that how these breaks are handled may play a role in the observed differential efficiency.

If these methylated vs. unmethylated inserts are handled differently, it might, but not necessarily, be reflected in a difference in the rates of repair-associated indels. We therefore examined the rates of indels at the flanking CRISPR/Cas9 cut sites, excluding sequences from 6-TG selected samples. We did not find a difference between the methylated and unmethylated alleles (48.9% vs. 50.9%, Fisher’s exact test: p ≈ 0.3). This is reflected in the similarity of the distributions of indel sizes for methylated vs. unmethylated sequences (**Fig. 3b**). In contrast, we observed a higher rate of indels for exogenous inserts (*i.e.* methylated and unmethylated alleles in either orientation) as compared with endogenous inserts (50.4% vs. 40.6%, Fisher’s Exact test: p < 2.2 × 10^−16^; size distribution of events in **Fig. 3c**). This suggests that exogenous DNA might be more likely to be inserted if there is exonuclease chew back during the repair. This result is further supported by the indel distribution of 6-TG-selected methylated and unmethylated alleles, which showed many indels extending from the CRISPR/Cas9 cut sites into the interior of the CpG island (**Fig. 3a**). Again excluding 6-TG selected sequences, we observed higher rates of indels for forward-oriented wild-type alleles as compared to inverted wild-type alleles (46.8% vs. 27.5%, Fisher’s exact test: p < 2.2 × 10^−16^; size distribution of events in **Fig. 3d**). However, this could simply be due to an increased propensity for indels when the break-repair recreates the wild-type sequence without mutation, because this site becomes a substrate for CRISPR/Cas9 cleavage again. This break-repair cycle can repeat until Cas9 is no longer active or a mutation occurs, explaining the higher rate of observed indels with forward-oriented wild-type alleles.

## Discussion

In this proof-of-concept study, we demonstrate concurrent epigenome and genome editing using CRISPR/Cas9. Our approach was to swap out endogenous DNA for exogenous DNA that was *in vitro* methylated and furthermore harbored programmed sequence differences. Specifically, we excised the endogenous *HPRT1* CpG island DNA using dual, flanking CRISPR/Cas9 cuts in the presence of transfected, *in vitro* methylated, SNV containing, exogenous *HPRT1* CpG island DNA. Our results demonstrate that it is possible to directly introduce *in vitro* methylated DNA into the genome using the NHEJ repair machinery in a targeted manner, and critically, that methylation of the exogenous fragment is maintained and can lead to robust gene silencing.

For targeted methylation, this CRISPR/NHEJ approach represents an alternative to the previously demonstrated dCas9-methyltransferase domain fusion protein approach (Liu et al. 2016; Amabile et al. 2016; Vojta et al. 2016; McDonald et al. 2016; Stepper et al. 2017; Lei et al. 2017; Xiong et al. 2017). While both approaches can produce targeted, scarless methylation of genomic DNA, the CRISPR/NHEJ approach is distinguished by the potential to program precisely which subsets of CpG dinucleotides are methylated, *e.g.* if exogenous inserts with specific patterns of CpG methylation are synthesized. In principle, this CRISPR/NHEJ strategy could be used to investigate the functional consequences of methylation patterns at single-site resolution, *e.g.* whether specific CpGs or combinations of CpGs are more important than others, and also whether/how these functional consequences depend on local sequence variation. Furthermore, other base modifications, *e.g.* hydroxymethylation or even non-standard bases, could be introduced into the genome by our approach, perhaps to study how they would be repaired or themselves further modified over subsequent cycles of DNA replication.

At least to our knowledge, this level of resolution is not possible with the dCas9-methyltransferase approach, which non-uniformly methylates sites over a window that may include tens to hundreds of CpGs in a probabilistic manner that depends on proximity to the enzyme (Liu et al. 2016; Amabile et al. 2016; Vojta et al. 2016; McDonald et al. 2016; Stepper et al. 2017; Lei et al. 2017; Xiong et al. 2017). Beyond resolution, a further advantage of the CRISPR/NHEJ approach is that it separates the effect of the methylated base from the act of methylation, *i.e.* functional effects observed with the dCas9-methyltransferase may be due to the effects of the fusion protein binding to the CpG island or promoter, rather than the methylated CpGs themselves.

These advantages notwithstanding, there are important practical limitations of our approach. There were three key elements of the experimental design that made this approach successful on the CpG island of *HPRT1*. First, rather than using RNA sequencing as a readout, we used selection for gene silencing and PacBio long-read DNA sequencing as a functional readout. This was necessary because of the diversity of editing outcomes and the fact that the vast majority did not involve the methylated allele (**Fig. 2a**; **Fig. 3a**). Second, since selection was required, we chose to target methylation to the *HPRT1* CpG Island. Expression of this gene in the presence of a small molecule chemotherapeutic, 6-TG, results in cell death. This allowed us to enrich for cells in which *HPRT1* had been successfully silenced. Third, we performed our experiments in the Hap1 cell line because it is haploid, such that the phenotype caused by successful insertion of the methylated allele would not be obscured by an unedited, expressed second copy of *HPRT1*, as would be the case with a diploid cell line.

In other experiments, we attempted to apply the CRISPR/NHEJ approach to methylate the CpG island of other genes. However, this proved difficult because of the requirement for a selection-based readout. Towards making such a readout possible on other genes beyond *HPRT1*, we engineered derivative Hap1 cell lines in which target genes were tagged with a negative selection marker such that expression of the gene would result in sensitivity to a small molecule drug, replicating the interaction between 6-TG and the *HPRT1* gene. Unfortunately, we were unable to successfully complete these experiments because of the poor transfection efficiency of the engineered cell lines. Freshly thawed, low passage HAP1 cells have a transfection efficiency of <5%, and after the many passages required for engineering, this reduced to approximately 0.1%. This low transfection efficiency is compounded by the low rate of NHEJ repair in Hap1 cells. Future studies employing this approach of tagging other genes with negative selection markers will need to use much larger numbers of Hap1 cells or alternative cell lines with similar properties to Hap1 cells, but with better transfection efficiencies.

Finally, an important limitation of our approach is the effectively low efficiency of methylation. This study showed much lower methylation rates (<1%) as compared to the dCas9-methyltransferase fusion protein approach (30-70%) (Liu et al. 2016; Amabile et al. 2016; Vojta et al. 2016; McDonald et al. 2016; Stepper et al. 2017; Lei et al. 2017; Xiong et al. 2017). This is due to a combination of factors, including the low transfection efficiency and rate of NHEJ of the Hap1 cell line, the lower integration rate of methylated DNA, and the availability of alternative outcomes that are also selected for, *e.g.* most prominently reinsertion of the endogenous DNA fragment in an inverted orientation. These limitations are potentially addressable through further modifications of the approach.

In conclusion, in this proof-of-concept study, we demonstrated concurrent epigenome and genome editing of the *HPRT1* CpG island in a single event using dual CRISPR/Cas9 cuts. The direct replacement of the native *HPRT1* CpG island sequence with the methylated exogenous *HPRT1* CpG island sequence resulted in functional *HPRT1* gene silencing. This approach constitutes a highly programmable new method for studying the direct effects of methylated DNA sequences in their endogenous contexts that may prove broadly useful for understanding the interplay between DNA modifications and gene expression at high resolution.

## Materials and Methods

### Generation of HPRT1 CpG Island Alleles and Guide RNAs

The HPRT1 CpG island region (GRCh37/hg19, chrX:133593033-133595157; **Supplemental Fig. 5**) was amplified from HeLa S3 DNA using Kapa Hifi Hotstart Readymix (Kapa Biosciences) and primers 1 and 2. Sequences of all primer and oligonucleotides used are in Supplemental Table 1. This amplicon was cloned using ClonTech In-Fusion Cloning kit into the pUC19 vector supplied with the kit. Synonymous SNVs were introduced into the cloned HPRT1 CpG island plasmid by PCR amplification of the entire plasmid with primers 3-6 using Kapa Hifi Hotstart Readymix (Kapa Biosciences) followed by re-circularization of the plasmid using ClonTech In-Fusion Cloning Kit. The synonymous SNVs were placed within the exon 1 coding sequence at genome positions, chrX:133594350 (C to T; Allele 1), chrX:133594353 (C to G; Allele 2), chrX:133594356 (C to T; Allele 2), and chrX:133594359 (T to A; Allele 1). For gRNAs, oligonucleotides 7-10 were synthesized by IDT, annealed, and cloned into pX458 plasmid (Addgene plasmid #48138) using ClonTech In-Fusion Cloning kit. The spacer sequences for these gRNAs were from chrX:133593802-133593821 and chrX:133594918-133594938. All cloned sequences were verified by Sanger Sequencing. DNA was extracted for all constructs using Qiagen mini-prep kits following manufacturer’s instructions on multiple 5 mL cultures.

To generate NHEJ template DNA, cloned alleles were amplified using Kapa Hifi Hotstart Readymix (Kapa Biosciences) and primers 11 and 12 resulting in an amplicon with the same sequence as chrX:133593819-133594938. This sequence is the region expected to be cut out of the genome by the gRNAs cloned above. The primers contain three phosphorothioate linkages at the 5’ end and mutations to destroy protospacer adjacent motif (PAM) sites at genome positions, chrX:133593824 (G to C) and chrX133594933 (C to G). PCR purification was performed using PCR Purification kit (Qiagen). DNA was methylated *in vitro* using M.SssI methyltransferase (NEB) following manufacturer’s instructions. To confirm methylation, DNA was digested using the methylation sensitive restriction enzyme, SmaI (NEB), following manufacturer’s instructions and visualized by polyacrylamide gel (SeaKem LE Agarose, Lonza) and SYBR Gold (Invitrogen). Methylated DNA was cleaned-up using a Qiagen PCR Purification kit. All concentrations were determined by using Qubit dsDNA BR kit (Invitrogen).

### Cell Culture, Transfections, FACS, and Selection

The haploid cell line Hap1 was maintained at 37°C in Iscove’s Modified Dulbecco’s Medium (ThermoFisher Scientific) supplemented with 10% fetal bovine serum and penicillin/streptomycin. For transfections, cells were trypsinized with 0.05% Trypsin-EDTA (ThermoFisher Scientifc) and reseeded in 10 cm dishes to achieve approximately 50% confluency by the next day. The next day each plate of cells was transfected with a mixture of both gRNA plasmids and both allele amplicons in a 0.45:0.45:0.05:0.05 ratio with a total of 18 ug of DNA per plate using Turbofectin 8.0 (Origene) and otherwise following manufacturer’s instructions. For 3 plates, the allele 1 template was methylated and the allele 2 template was unmethylated. For the other 3 plates, the allele 2 template was methylated and allele 1 template was unmethylated. Forty-eight hours after transfection, cells were trypsinized and incubated for 45 minutes at 37°C in medium containing 10 ug/mL Hoechst 33342 (ThermoFisher Scientific), a live-cell DNA dye. Fluorescence Activated Cell Sorting (FACS) was used to retrieve more than 100,000 cells from each plate that were both GFP positive (i.e., transfected) and in the G1 cell cycle phase (i.e., haploid). Sorted cells were placed back into culture in 6-well dishes for 1 week in supplemented medium with media changes every 3 days. At 1 week, each dish of cells was trypsinized and washed with Dulbecco’s phosphate buffered saline (ThermoFisher Scientific). Fifty percent of each sample of cells was snap frozen for later DNA extraction and the other fifty percent was split to two wells of a 6-well dish. One of these wells received 5 µM 6-TG (Sigma) in DMSO for negative selection and the other received DMSO as a control (mock selection). A control plate of untransfected cells was also treated with 5 µM 6-TG to monitor selection status. Cells were cultured for 11 days with media changes and replacement of selection agents every 3 days. At 11 days, cell were trypsinized and snap frozen for later DNA extraction.

### DNA Extraction and Sequencing

DNA and RNA were extracted using a Qiagen Allprep kit as per manufacturer’s instructions. For Illumina sequencing, a three-round nested PCR with Kapa Hifi Hotstart Readymix and 250 ng of DNA (~100,000 genome equivalents) per sample was used to prepare amplicons. The first round of PCR with 3 cycles (primers 13 and 14) added a unique molecular index (UMI), the second round (primers 15 and 16) was for amplification, and the third round (primers 17-27) added flow-cell adapters starting with 1/50 of the second-round reaction as input. Round 2 and 3 PCRs were followed in real time using SYBR Green (Invitrogen) and stopped prior to plateauing. Agencourt Ampure XP bead (Beckman-Coulter) clean-up (1.0X) was performed after each round of PCR. Amplicon DNA from each sample was pooled at equal concentration and sequenced on an Illumina MiSeq using a 2 × 75 cycle paired-end kit with custom sequencing (primers 51 and 52) and index primers (primer 53), but otherwise as per manufacturer’s instructions.

For Pacific Biosciences sequencing, a two-round nested PCR with Kapa Hifi Hotstart Readymix, and 250 ng of DNA per sample was used to prepare amplicons. The first round with 3 cycles added a UMI to some of the samples (primers 28 and 29) or added a UMI and sample barcode to the remaining samples (primers 29 and 32-45), and the second round (primers 30 and 31) was for amplification starting with 1/50 of the second-round reaction as input. To increase the amount of DNA prior to gel extraction for samples without barcodes, a third round of PCR using the second-round primers was performed. Round 2 and 3 PCRs were followed in real time using SYBR Green (Invitrogen) and stopped prior to plateauing. Using SYBR Gold and blue light for visualization, gel extractions of the approximately 2000 bp band were performed to reduce the number of deletions (approximately 1000 bp) sequenced. For samples without barcodes, different 1.5% agarose gels were used for each sample. For barcoded samples, groups of samples were pooled prior to loading the gel and groups of pools were gel extracted together. A Qiagen Gel Extraction Kit was used as per manufacturer’s instructions. For samples without barcodes, 500 ng of DNA per sample were used as input into Pacific Biosciences SMRT Bell Template Prep Kit 1.0 for preparation for sequencing as per manufacturer instructions. For samples with barcodes, the gel extracted DNA pools were mixed at equal concentrations and then prepared for sequencing by the University of Washington PacBio Sequencing Service (UWPBSS). For samples without barcodes, sequencing was performed on an RSII using P6-C4 chemistry by the UWPBSS using one SMRT cell per sample. For samples with barcodes, the library was sequenced on a Sequel SMRT Cell 1M v3.0.

For bisulfite sequencing, between 420 ng and 1344 ng of DNA per sample was bisulfite converted using Promega MethylEdge Bisulfite Converion kit as per manufacturer’s instructions. A 3 round nested PCR with Kapa Hifi Uracil+ (first and second rounds) and Kapa Hifi Hotstart Readymix (third round) and half of the bisulfite converted DNA was used to prepare amplicons for Illumina sequencing. The first round was 3 cycles (primers 46 and 47) for adding UMIs, the second round (primers 48 and 49) was for amplification, and third round (primers 17-24 and 50) was for adding flow-cell adapters starting with 1/50 of the second-round reaction as input. Round 2 and 3 PCRs were followed in real time and stopped prior to plateauing. Agencourt Ampure XP bead clean-ups (0.8X) were performed twice after each round of PCR. Amplicon DNA from each sample was pooled and sequenced on a MiSeq using 2 × 250 cycle paired-end kit with custom sequencing and index primers (primers 51-53).

### Sequencing Data Analysis

For Illumina DNA sequencing, after bcl2fastq (version 2.18, Illumina) was run for demultiplexing, read 2 FASTQ files were converted to FASTA format. Sequences were then converted to their reverse complement and aligned to the HPRT1 CpG island reference (chrX:133594298-133594522) sequence using needleall (version EMBOSS:6.5.7.0, http://emboss.sourceforge.net/apps/release/6.5/emboss/apps/needleall.html). Based on this alignment, sequences were assigned to alleles (allele 1 vs. allele 2 vs. wild type) using allele defining SNVs. Perfect matches of all bases in a portion of exon 1 (chrX:133589685-133600894) including the coding sequence and at the 4 SNV positions were required for assignment to an allele group.

For bisulfite sequencing, after bcl2fastq was run for demultiplexing, paired-end reads were merged with PEAR (Paired-End reAd mergeR, version 0.9.6) and discordant pairs were removed (Jiang et al. 2014). Sequences were then converted to their reverse complement and aligned using needleall to the HPRT1 CpG island reference (chrX:133594296-133594578) sequences consisting of a bisulfite converted sequence, a bisulfite converted sequence assuming all CpGs were methylated, and an unconverted sequence. UMIs and HPRT1 CpG island sequences were extracted from the BAM files for each read based on the alignment. The sequences were clustered by UMI and a consensus sequence was generated for each cluster by simple majority at each position in the sequence. The consensus sequences were then realigned with the reference sequences using needleall. Based on this alignment, sequences were assigned to alleles (allele 1 vs. allele 2 vs. wild type) using the allele-defining SNVs. Perfect matches of all bases in a portion of exon 1, including the coding sequence, (chrX:133594296-133594578) and at the 4 SNV positions were required for assignment to an allele group.

For Pacific Biosciences sequencing data, bax2bam (version 0.0.2, Pacific Biosciences, Inc.) was run on the .h5 files for conversion to BAM files. This was followed by circular consensus calling using CCS (version 2.0.0, Pacific Biosciences, Inc.). Sequences from the generated BAM files were converted to their reverse complement and both the forward and reverse complement sequences were saved to FASTA format. All sequences were aligned using needleall against the reference forward and inverted sequences of the HPRT1 CpG island. Reference sequences included the HPRT1 CpG island sequence and flanking primer sequences to allow UMIs to be captured. Barcodes were also included in the reference sequences for the Sequel SMRT cell sequencing data to assign each read to a sample. The inverted reference was created by inverting the sequence between the CRISPR cut sites, but keeping the flanking sequence unaltered. The UMIs and HPRT1 CpG island sequences were extracted from the BAM alignment files for each read based on alignment coordinates. Again, sequences were clustered by UMI, a consensus sequence calculated and realigned using needleall. Based on this new alignment, sequences were grouped by alleles (allele 1 vs. allele 2 vs. wild type vs. deletion) and orientation (forward vs. inverted) using the 4 allele defining SNVs and 2 PAM mutations. Perfect matches in the promoter, exon 1, and splice donor sequence (chrX:133594123-133594374), and at the allele-defining SNVs and PAM positions were required for assignment to an allele group.

Counts of reads assigned to allele groups were used for **Fig. 2**, as described in the figure caption. For **Fig. 3**, indels were counted in reads assigned to allele groups. Specifically, indels within 5 bp on either side of the expected CRISPR/Cas9 cut sites based on the read alignments above were included in the count. Sizes of these indels were also determined based on the alignment. The deletion could only extend 4 bases into the insert sequence because the PAM mutation which was at the fifth base was required for assignment to an allele group. Unless otherwise noted, custom scripts were written for this analysis using bash, Python, or R programming languages.

## Acknowledgments

Jason Klein and Seungsoo Kim provided helpful comments. Donna Prunkard and the University of Washington Pathology Flow Cytometry Core Facility were very helpful with FACS.

## Declarations

The datasets generated and/or analysed during the current study are available in the SRA repository, http://www.ncbi.nlm.nih.gov/bioproject/547358

## Supplemental Figures

**Supplemental Figure 1:**
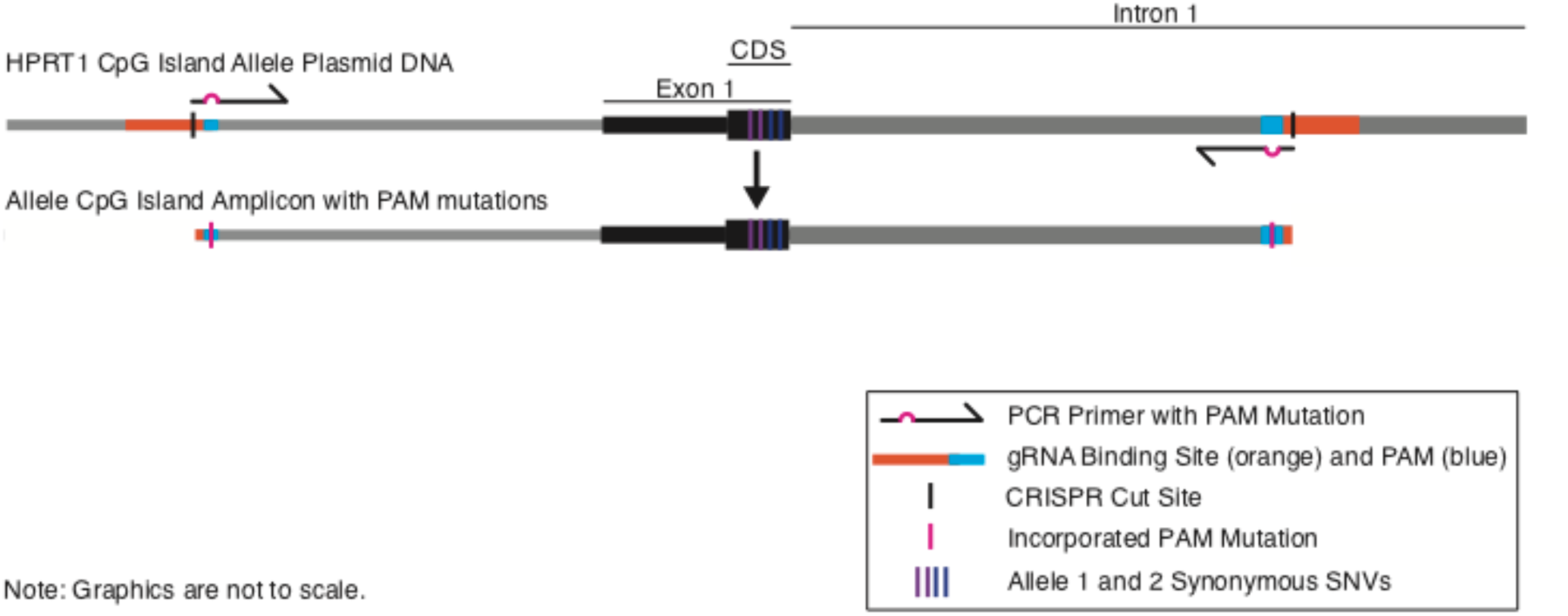
PCR for generation of CpG Island Allele Amplicon DNA with PAM mutations. Cloned *HPRT1* CpG island plasmid DNA was used as a template for PCR amplification. PAM sites corresponding to the guide RNA target sites are at the ends within the CpG island amplicon. Primer sequences included a mismatch near the 5’ end resulting in incorporation of a mutation in the PAM sequences at the ends of the CpG island allele amplicons.

**Supplemental Figure 2:**
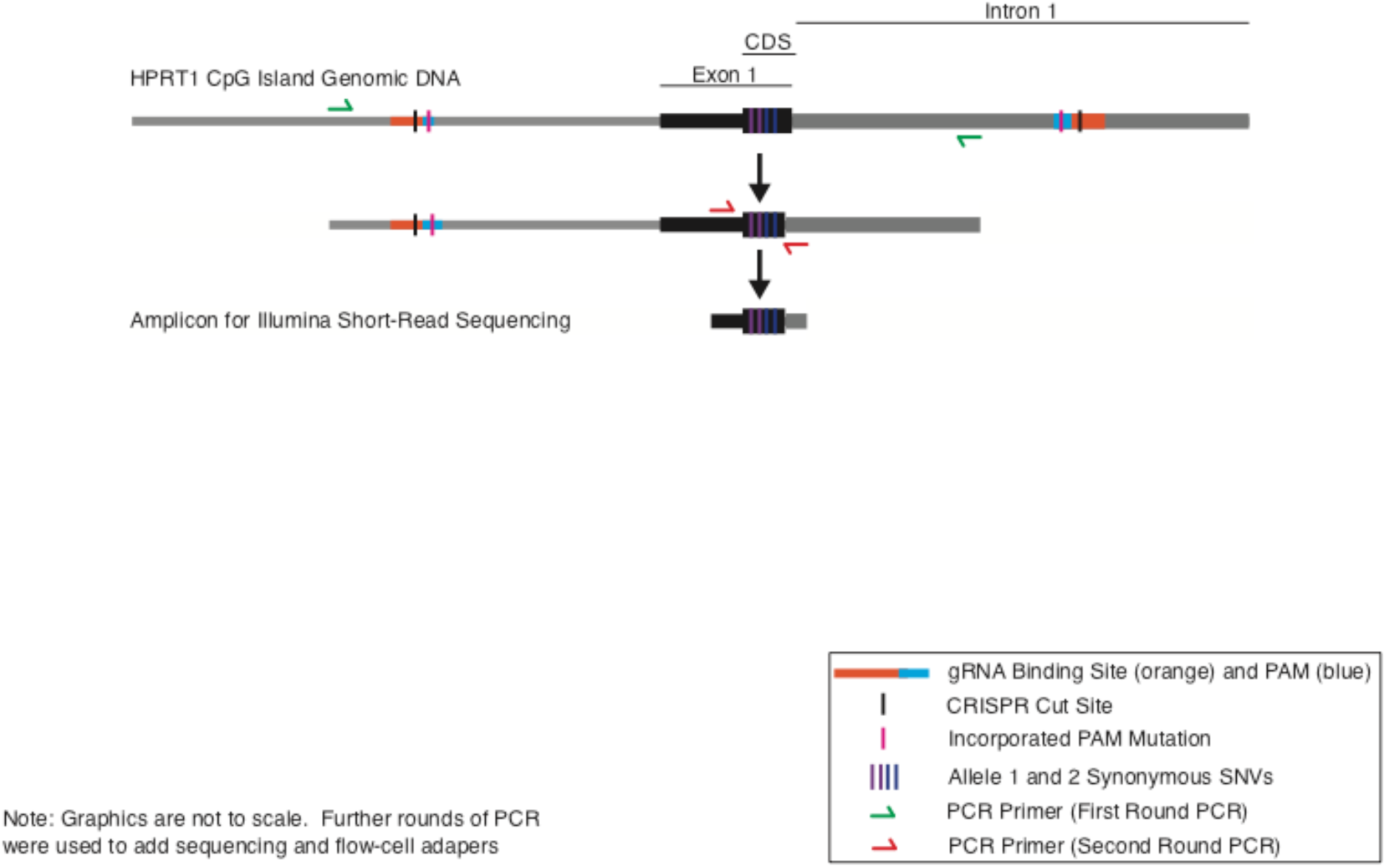
Nested PCR from Pre-selection, Mock Selection, or 6-TG Selected Cell Genomic DNA for Illumina Sequencing. Genomic DNA was the template for the first round of PCR. In this round, one primer was outside the CRISPR cut sites in the genome, while the other primer was within the cut sites in the *HPRT1* CpG island. The product of the PCR was used as the template for the second round PCR. The second round PCR amplified a 44-bp region including the allele-defining SNVs.

**Supplemental Figure 3:**
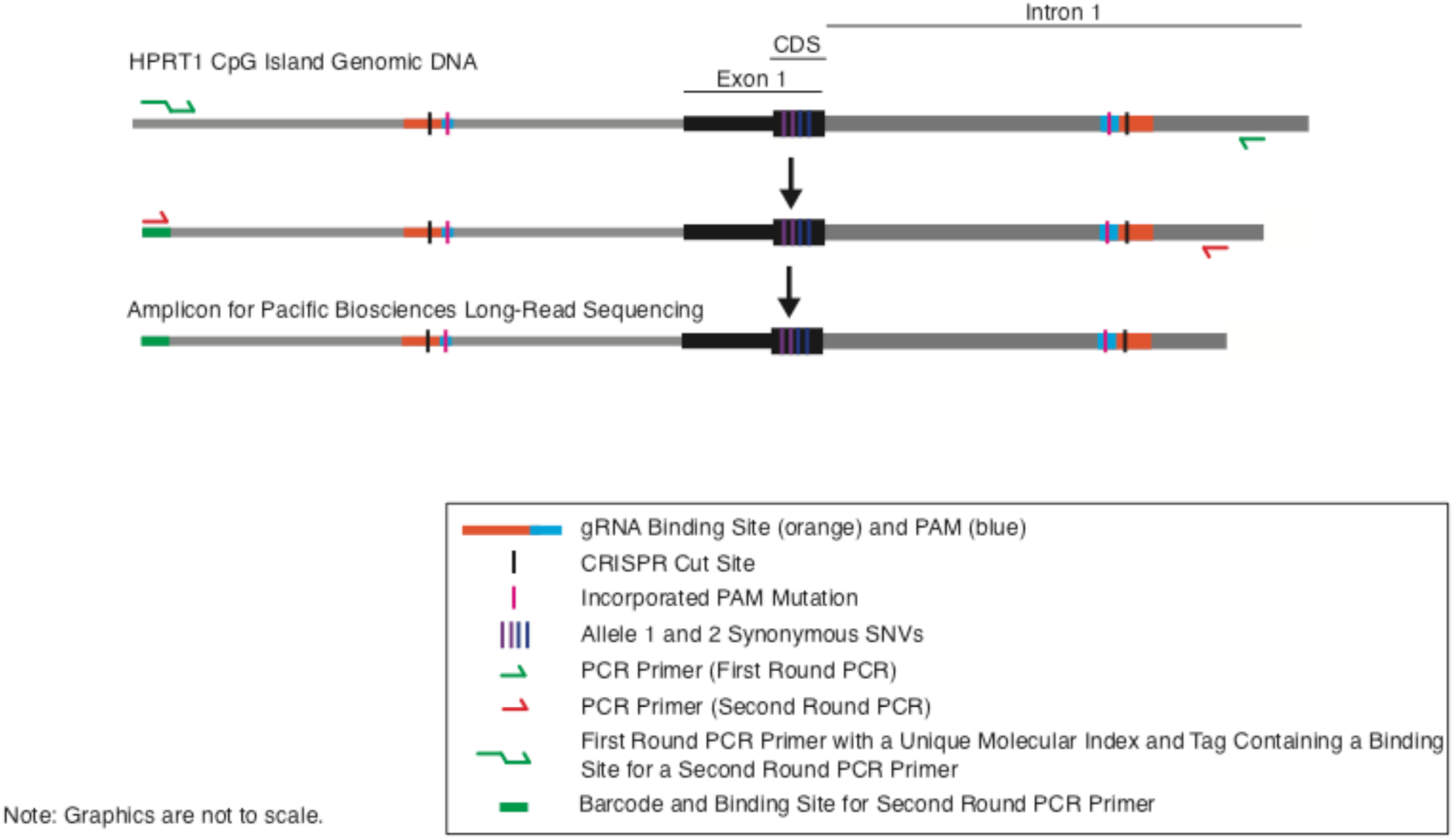
PCR from Pre-selection, Mock Selection, or 6-TG Selected Cell Genomic DNA for Pacific Biosciences Sequencing. Genomic DNA was the template for the first round of PCR. In this round, both primers were outside the CRISPR cut sites in the genome. An unique molecular index (UMI) and a primer binding site were added in this round of PCR. The product of this PCR was used as the template for the second round PCR.

**Supplemental Figure 4:**
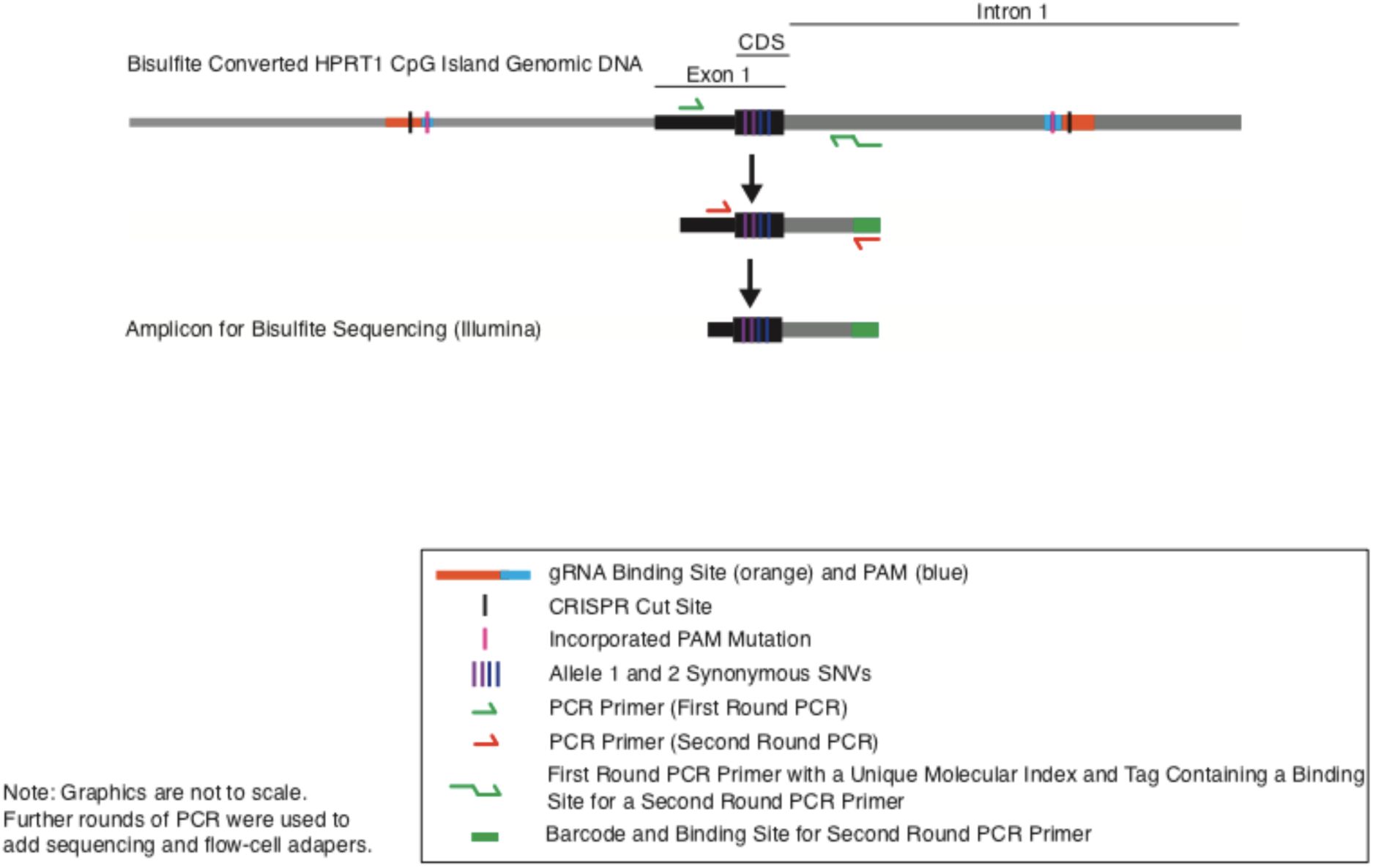
Nested PCR from Bisulfite-Converted Pre-selection, Mock Selection, or 6-TG Selected Cell Genomic DNA for Illumina Sequencing. Bisulfite-converted genomic DNA was the template for the first round of PCR. In this round, both primers were inside the CRISPR cut sites in the genome. An unique molecular index (UMI) and a primer binding site were added in this round of PCR. The product of this PCR was used as the template for the second round PCR.

**Supplemental Figure 5.**
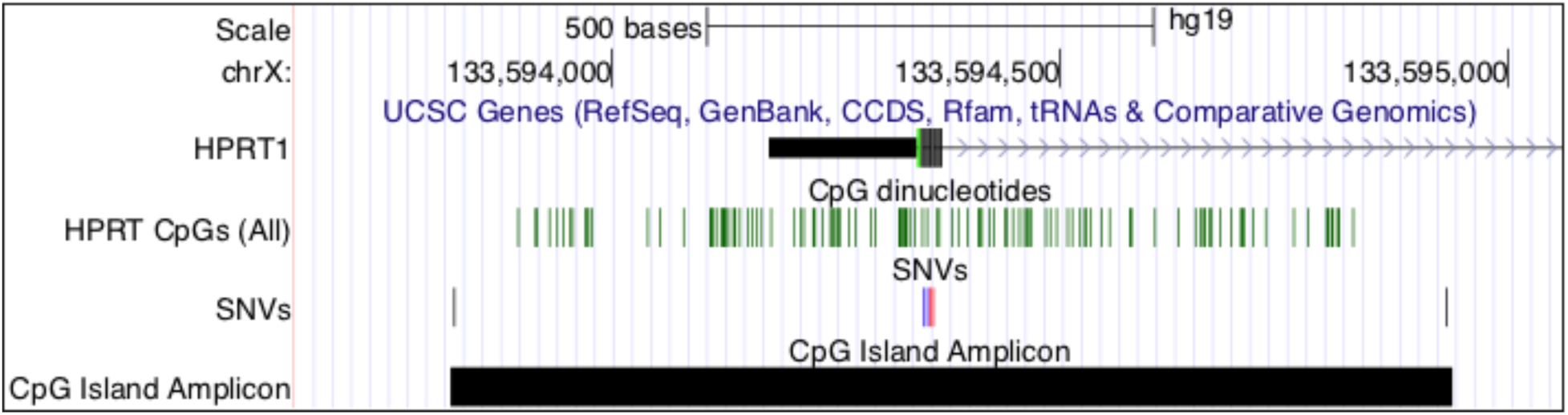
UCSC genome browser view showing a region around the transcriptional start of *HPRT1*, CpG dinucleotides included in the CpG island amplicons, locations of SNVs introduced to create alleles and destroy PAM sites, and location of the *HPRT1* CpG island.

## Supplemental Table 1: Primers

### HPRT1 CpG island region genomic amplification

1. (JA-Me-HPRT-gen-1-f): CGG TAC CCG GGG ATC-gtgaggcaaaaatagaggctcagagt

2. (JA-ME-HPRT-gen-1-R): CGA CTC TAG AGG ATC GCA GCT TGG CCG GTT CAA CAAA

### Synonymous SNVs

#### Allele 1 (SNVs at 133594350 and 133594359)

3. Forward (JA-Me-HPRT-Mut-6-f): T-CGCAGCCC-A-GGCGT + CGTGGTGAGCAGCTCGGC

4. Reverse (JA-Me-HPRT-Mut-6-r): ACG CCT GGG CTG CGA + GTCGCCATAACGGAGCCG

#### Allele 2 (SNVs at 133594353 and 133594356)

5. Forward (JA-Me-HPRT-Mut-7-f): CCG-G-AG-T-CCTGGCGT + CGTGGTGAGCAGCTCGGC

6. Reverse (JA-Me-HPRT-Mut-7-r): ACG CCA GGA CTC CGG + GTCGCCATAACGGAGCCG

### Oligos for cloning gRNAs

#### chrX:133593694-133595157

7. (JA-Me-HPRT-sgRNA5’-1-f) CACC-GCAACTGTTACAACCAGTTAA

8. (JA-Me-HPRT-sgRNA5’-1-r) aaac-TTAACTGGTTGTAACAGTTG-C

#### chrX:133593694-133595157

9. (JA-Me-HPRT-sgRNA3’-1-f) CACCGcttcgtgtgtcaaatacgca

10. (JA-Me-HPRT-sgRNA3’-1-r) AAAC-tgcgtatttgacacacgaag-C

### NHEJ Template Amplicon

11. (JA-ME-HPRT-Nokin-1*-f): T*A*A*GG-C-TTTGGGGAAGCACTG

12. (JA-ME-HPRT-Nokin-1*-r): g*c*a*tg-c-ttaccgctaccagag

* = phosphorothioate linkage

### Illumina Sequencing

#### Round 1

13. (JA-ME-HPRT-onDNA-1-f): GTGAGGCAAAAATAGAGGCTCAGAGT

14. (JA-ME-HPRT-onDNA-2-r): CAGTGTGTGCAAAACTAAAGGCA

#### Round 2

15. (JA-ME-HPRT-cDNA-1-f): CTAAATGGCTGTGAGAGAGCTCAG-TTCCTCCTCCTGAGCAGTCAGC

16. (JA-ME-HPRT-DNA-1-r): ACTTTATCAATCTCGCTCCAAACC-CGAGCTGCTCACCACGA

#### Round 3 Reverse

17. (jkA0142_ampRbnew1_codv): CAAGCAGAAGACGGCATACGAGAT AAGCGTTCA gaccgtcggc ACTTTATCAATCTCGCTCCAAACC

18. (jkA0143_ampRbnew2_codv): CAAGCAGAAGACGGCATACGAGAT CGCAAGCGT gaccgtcggc ACTTTATCAATCTCGCTCCAAACC

19. (jkA0144_ampRbnew3_codv): CAAGCAGAAGACGGCATACGAGAT GCAGCGCGA gaccgtcggc ACTTTATCAATCTCGCTCCAAACC

20. (jkA0145_ampRbnew4_codv): CAAGCAGAAGACGGCATACGAGAT CGCGCAGCT gaccgtcggc ACTTTATCAATCTCGCTCCAAACC

21. (jkA0146_ampRbnew5_codv): CAAGCAGAAGACGGCATACGAGAT TCAAGCGCA gaccgtcggc ACTTTATCAATCTCGCTCCAAACC

22. (jkA0147_ampRbnew6_codv): CAAGCAGAAGACGGCATACGAGAT CAGTCGCAG gaccgtcggc ACTTTATCAATCTCGCTCCAAACC

23. (jkA0148_ampRbnew7_codv): CAAGCAGAAGACGGCATACGAGAT GCGTCAGTT gaccgtcggc ACTTTATCAATCTCGCTCCAAACC

24. (jkA0149_ampRbnew8_codv): CAAGCAGAAGACGGCATACGAGAT AGTCGCGCA gaccgtcggc ACTTTATCAATCTCGCTCCAAACC

#### Round 3 Forward

25. (P5_pu1Lfwd_fwd_GF_P5barc_03): AATGATACGGCGACCACCGAGATCTACACAAATCTGCGT acgtaggc CTAAATGGCTGTGAGAGAGCTCAG

26. (P5_pu1Lfwd_fwd_GF_P5barc_04): AATGATACGGCGACCACCGAGATCTACACATTTAGTACG acgtaggc CTAAATGGCTGTGAGAGAGCTCAG

27. (P5_pu1Lfwd_fwd_GF_P5barc_05): AATGATACGGCGACCACCGAGATCTACACCCACACAAGC acgtaggc CTAAATGGCTGTGAGAGAGCTCAG

### Pacific Biosciences RSII Sequencing

#### Round 1

28. (JA-MeHPRT-InDel-Out1-f): CTAAATGGCTGTGAGAGAGCTCAG-NNNNN NNNNN-AATGGTGTTGCTGGAGCAACTGTT

29. (JA-MeHPRT-InDel-Out1-r): GCAGGCTAAAGCATATTTAACTGGC

#### Round 2 and 3

30. (JA-MeHPRT-InDel-In1-f): GGTGGT-GAATTC-CTAAATGGCTGTGAGAGAGCTCAG

31. (JA-MeHPRT-InDel-In1-r): GGTGGT-CCTGCAGG-GTGGAGCTAAGATACCAGAGGCTG

### Pacific Biosciences Sequel Sequencing

#### Round 1 Forward

32. (JA-MeHPRT-142-Out1-f): CTAAATGGCTGTGAGAGAGCTCAG-TGAACGCTT-NNNNN NNNNN-AATGGTGTTGCTGGAGCAACTGTT

33. (JA-MeHPRT-143-Out1-f): CTAAATGGCTGTGAGAGAGCTCAG-ACGCTTGCG-NNNNNNNNNN-AATGGTGTTGCTGGAGCAACTGTT

34. (JA-MeHPRT-144-Out1-f): CTAAATGGCTGTGAGAGAGCTCAG-TCGCGCTGC-NNNNNNNNNN-AATGGTGTTGCTGGAGCAACTGTT

35. (JA-MeHPRT-145-Out1-f): CTAAATGGCTGTGAGAGAGCTCAG-AGCTGCGCG-NNNNNNNNNN-AATGGTGTTGCTGGAGCAACTGTT

36. (JA-MeHPRT-146-Out1-f): CTAAATGGCTGTGAGAGAGCTCAG-TGCGCTTGA-NNNNNNNNNN-AATGGTGTTGCTGGAGCAACTGTT

37. (JA-MeHPRT-147-Out1-f): CTAAATGGCTGTGAGAGAGCTCAG-CTGCGACTG-NNNNN NNNNN-AATGGTGTTGCTGGAGCAACTGTT

38. (JA-MeHPRT-InDel_148-Out1-f): CTAAATGGCTGTGAGAGAGCTCAG-AACTGACGC-NNNNNNNNNN-AATGGTGTTGCTGGAGCAACTGTT

39. (JA-MeHPRT-InDel_149-Out1-f): CTAAATGGCTGTGAGAGAGCTCAG-TGCGCGACT-NNNNNNNNNN-AATGGTGTTGCTGGAGCAACTGTT

40. (JA-MeHPRT-InDel_151-Out1-f): CTAAATGGCTGTGAGAGAGCTCAG-ACATCGTAG-NNNNNNNNNN-AATGGTGTTGCTGGAGCAACTGTT

41. (JA-MeHPRT-InDel_152-Out1-f): CTAAATGGCTGTGAGAGAGCTCAG-TAGCAGGAT-NNNNN NNNNN-AATGGTGTTGCTGGAGCAACTGTT

42. (JA-MeHPRT-InDel_153-Out1-f): CTAAATGGCTGTGAGAGAGCTCAG-ATCGTAGCC-NNNNNNNNNN-AATGGTGTTGCTGGAGCAACTGTT

43. (JA-MeHPRT-InDel_154-Out1-f): CTAAATGGCTGTGAGAGAGCTCAG-TCTATTCAC-NNNNNNNNNN-AATGGTGTTGCTGGAGCAACTGTT

44. (JA-MeHPRT-InDel_155-Out1-f): CTAAATGGCTGTGAGAGAGCTCAG-CGTCTCCTA-NNNNNNNNNN-AATGGTGTTGCTGGAGCAACTGTT

45. (JA-MeHPRT-InDel_156-Out1-f): CTAAATGGCTGTGAGAGAGCTCAG-AGGTGATTC-NNNNNNNNNN-AATGGTGTTGCTGGAGCAACTGTT

#### Round 1 Reverse

29. (JA-MeHPRT-InDel-Out1-r): GCAGGCTAAAGCATATTTAACTGGC

#### Round 2 and 3

30. (JA-MeHPRT-InDel-In1-f): GGTGGT-GAATTC-CTAAATGGCTGTGAGAGAGCTCAG

31. (JA-MeHPRT-InDel-In1-r): GGTGGT-CCTGCAGG-GTGGAGCTAAGATACCAGAGGCTG

### Bisulfite Sequencing

#### Round 1

46. (JA-ME-HPRT-BCOutIll-2-f): CTAAATGGCTGTGAGAGAGCTCAG-NNNNN NNNNN-AGGAGGGAtttAttttAAAttt

47. (JA-ME-HPRT-BCOutIll-2-r): CCTCCTCCTCTaCTCc

#### Round 2

48. (JA-ME-HPRT-BCInIll-2-f): CTAAATGGCTGTGAGAGAGCTCAG

49. (JA-ME-HPRT-BCInIll-2-r): ACTTTATCAATCTCGCTCCAAACC-aCTTCCTCCTCCTaAaCAaTCAaCC

#### Round 3 Reverse

17. (jkA0142_ampRbnew1_codv): CAAGCAGAAGACGGCATACGAGAT AAGCGTTCA gaccgtcggc ACTTTATCAATCTCGCTCCAAACC

18. (jkA0143_ampRbnew2_codv): CAAGCAGAAGACGGCATACGAGAT CGCAAGCGT gaccgtcggc ACTTTATCAATCTCGCTCCAAACC

19. (jkA0144_ampRbnew3_codv): CAAGCAGAAGACGGCATACGAGAT GCAGCGCGA gaccgtcggc ACTTTATCAATCTCGCTCCAAACC

20. (jkA0145_ampRbnew4_codv): CAAGCAGAAGACGGCATACGAGAT CGCGCAGCT gaccgtcggc ACTTTATCAATCTCGCTCCAAACC

21. (jkA0146_ampRbnew5_codv): CAAGCAGAAGACGGCATACGAGAT TCAAGCGCA gaccgtcggc ACTTTATCAATCTCGCTCCAAACC

22. (jkA0147_ampRbnew6_codv): CAAGCAGAAGACGGCATACGAGAT CAGTCGCAG gaccgtcggc ACTTTATCAATCTCGCTCCAAACC

23. (jkA0148_ampRbnew7_codv): CAAGCAGAAGACGGCATACGAGAT GCGTCAGTT gaccgtcggc ACTTTATCAATCTCGCTCCAAACC

24. (jkA0149_ampRbnew8_codv): CAAGCAGAAGACGGCATACGAGAT AGTCGCGCA gaccgtcggc ACTTTATCAATCTCGCTCCAAACC

#### Round 3 Forward

50. (P5_pu1Lfwd_fwd_GF_P5barc_01): AATGATACGGCGACCACCGAGATCTACACTAAATGCTCC acgtaggc CTAAATGGCTGTGAGAGAGCTCAG

### Illumina Sequencing Primers

51. (PU1_SEQ_F_R1) acgtaggcCTAAATGGCTGTGAGAGAGCTCAG

52. (PU1_SEQ_R_R2) gaccgtcggcACTTTATCAATCTCGCTCCAAACC

53. (PU1_seqIx1) GGTTTGGAGCGAGATTGATAAAGTgccgacggtc

